# Rapid GMP-compliant expansion of SARS-CoV-2-specific T cells from convalescent donors for use as an allogeneic cell therapy for COVID-19

**DOI:** 10.1101/2020.08.05.237867

**Authors:** Rachel S Cooper, Alasdair R Fraser, Linda Smith, Paul Burgoyne, Stuart N Imlach, Lisa M Jarvis, Sharon Zahra, Marc L. Turner, John DM Campbell

**Author notes:** contributed equally to this work (Rachel S Cooper, Alasdair R Fraser). Corresponding author. Prof. John Campbell, Scottish National Blood Transfusion Service, The Jack Copland Centre, 52 Research Avenue North, Edinburgh, UK.

## Abstract

COVID-19 disease caused by the SARS-CoV-2 virus is characterized by dysregulation of effector T cells and accumulation of exhausted T cells. T cell responses to viruses can be corrected by adoptive cellular therapy using donor-derived virus-specific T cells. Here we show that SARS-CoV-2-exposed blood donations contain CD4 and CD8 memory T cells specific for SARS-CoV-2 spike, nucleocapsid and membrane antigens. These peptides can be used to isolate virus-specific T cells in a GMP-compliant process. These T cells can be rapidly expanded using GMP-compliant reagents for use as a therapeutic product. Memory and effector phenotypes are present in the selected virus-specific T cells, but our method rapidly expands the desirable central memory phenotype. A manufacturing yield ranging from 10^10^ to 10^11^ T cells can be obtained within 21 days culture. Thus, multiple therapeutic doses of virus-specific T cells can be rapidly generated from convalescent donors for treatment of COVID-19 patients

**One Sentence Summary:** CD4+ and CD8+ T cells specific for SARS-CoV-2 can be isolated from convalescent donors and rapidly expanded to therapeutic doses at GMP standard, maintaining the desired central memory phenotype required for protective immune responses against severe COVID-19 infections.

## Introduction

Coronavirus disease 2019 (COVID-19), caused by the severe acute respiratory virus syndrome – coronavirus 2 (SARS-CoV-2) emerged in Wuhan, China in December 2019. In the majority of cases infection with SARS-CoV-2 is asymptomatic or leads to relatively mild self-limiting disease, but a proportion of patients progress to severe disease with about a 1% overall mortality rate (1,2). Declared a pandemic by the WHO on 11^th^ March 2020, the virus has spread rapidly to all parts of the world with >15 million infections and >600,000 deaths reported by July 2020 (3).

Patients with progressive severe disease demonstrate a high neutrophil to lymphocyte ratio and a lymphopenia in the blood accompanied by a hyperinflammatory and prothrombotic diathesis leading to Acute Respiratory Distress Syndrome (ARDS) and multiorgan failure (4,5,6). Some success in treating severe disease has recently been reported with therapeutic agents such as remdesivir (7), dexamethasone (8), and nebulized interferon-beta (9).

A particular feature of progressive COVID-19 disease is rapid exhaustion of the memory T cell compartment – characterized by overall lymphopenia and accumulation of naïve/exhausted T cell memory phenotypes (10,11). This undesirable phenotype is associated with a systemic hyperinflammatory response and poor outcomes (reviewed in 12). Conversely, protection in self-limiting disease is associated with strong CD4 and CD8 T cells responses to the spike, membrane and nucleocapsid proteins of the virus and development of virus-specific antibodies (13–15). Convalescent plasma (CP) is currently being trialed in a number of countries as a potential therapeutic option, although the level and duration of protection afforded by the antibody response against re-infection remains unclear at present (16).

New therapies to support the immune response to SARS-CoV-2, preventing the collapse of the lymphocyte compartment and supporting protective immunity would have significant impact on outcome for hospitalized patients. Anti-viral T cells specific for viruses such as cytomegalovirus (CMV), adenovirus (ADV) and Epstein Barr Virus (EBV) have been successfully used as adoptive cellular therapies to combat such infections in patients with immune deficiency (17–22). Following selection of antigen-specific T cells from a blood donation from an individual who has been infected with the relevant virus, T cells may be expanded *in vitro* to manufacture banks of T cells from HLA-typed donors for repeated infusions in multiple patients with a partial HLA-match. We and others have adopted this approach in the treatment of EBV or CMV-driven disease (21, 22) with evidence of disease remission and low incidence of Graft versus Host Disease (GvHD) (21, 22).

In this study we present clear evidence to show that donations from individuals who have been infected with SARS-CoV-2 with mild symptoms and have recovered retain normal T cell compartment profiles, with CD4 and CD8 memory and effector T cells specific for SARS-CoV-2 spike, nucleocapsid and membrane antigens. These virus-specific T cells (VSTs) can be isolated using Good Manufacturing Practice (GMP)-compatible selection technology and rapidly expanded *in vitro* using closed culture vessels and GMP-compliant reagents and medium. The mononuclear cell fraction of a single whole blood donation from a COVID-19 convalescent donor (CCD) can be used to generate up to 10^11^ T cells within 21 days with the desired central memory phenotype as a potential new therapy for SARS-CoV-2. This offers the potential for the manufacture of a bank of HLA-matched donor T cell products for use in clinical trial and future treatment of COVID-19 patients.

## Results

### Donor characteristics and leucocyte phenotype

Buffy coats from CCD (n=15, see Table 1) were collected between 34-56 days after resolution of symptoms (diagnosis and resolution of infection were confirmed by SARS-CoV-2 PCR). Donors were 23-58 years old (median 49) and evenly split by gender (7 female, 8 male). In all cases, donors exhibited mild symptoms of COVID-19 infection and did not require hospital treatment. Immunophenotyping of buffy coat-isolated peripheral blood mononuclear cells (PBMC) from CCD compared to uninfected donors (UD, n=17) is shown in Figure 1. PBMC were sequentially gated as per Supplementary Figure 1. The mean percentage of T cells (CD3+/ CD56−), NKT cells (CD56+/ CD3+) and monocytes (CD14+) was comparable between UD and CCD (Fig 1A). The mean percentage of T cells with an activated phenotype (HLA-DR+/ CD38+), reported as elevated in other studies with moderate to severe disease, were not found to be significantly different between UD and CCD in this study. NK cell levels were significantly elevated (p=0.0073) in CCD compared to UD and the mean percentage of B cells in CCD was significantly lower than UD (p=0.0003). In this study, age did not correlate with NK cell or B cell levels in CCD (Fig 1B/C), though a significant correlation (Pearson correlation p=0.04, r=0.524) was identified between percentage of B cells and SARS-CoV-2 antibody content (Fig 1D). Within the T cell compartment the percentage of CD4 and CD8 T cells, as well as CD3+/CD4−/CD8− (double negative) and CD3+/CD4+/CD8+ (double positive) remained unchanged between CCD and UD (Figure 1E). In addition, analysis of co-expression of T cell memory markers CD62L, CD45RO and CD45RA reveals no difference in CD4 and CD8 memory subpopulations between UD and CCD for either CD4 T cells (Fig 1F) or CD8 T cells (Figure 1G).

**Figure 1.**
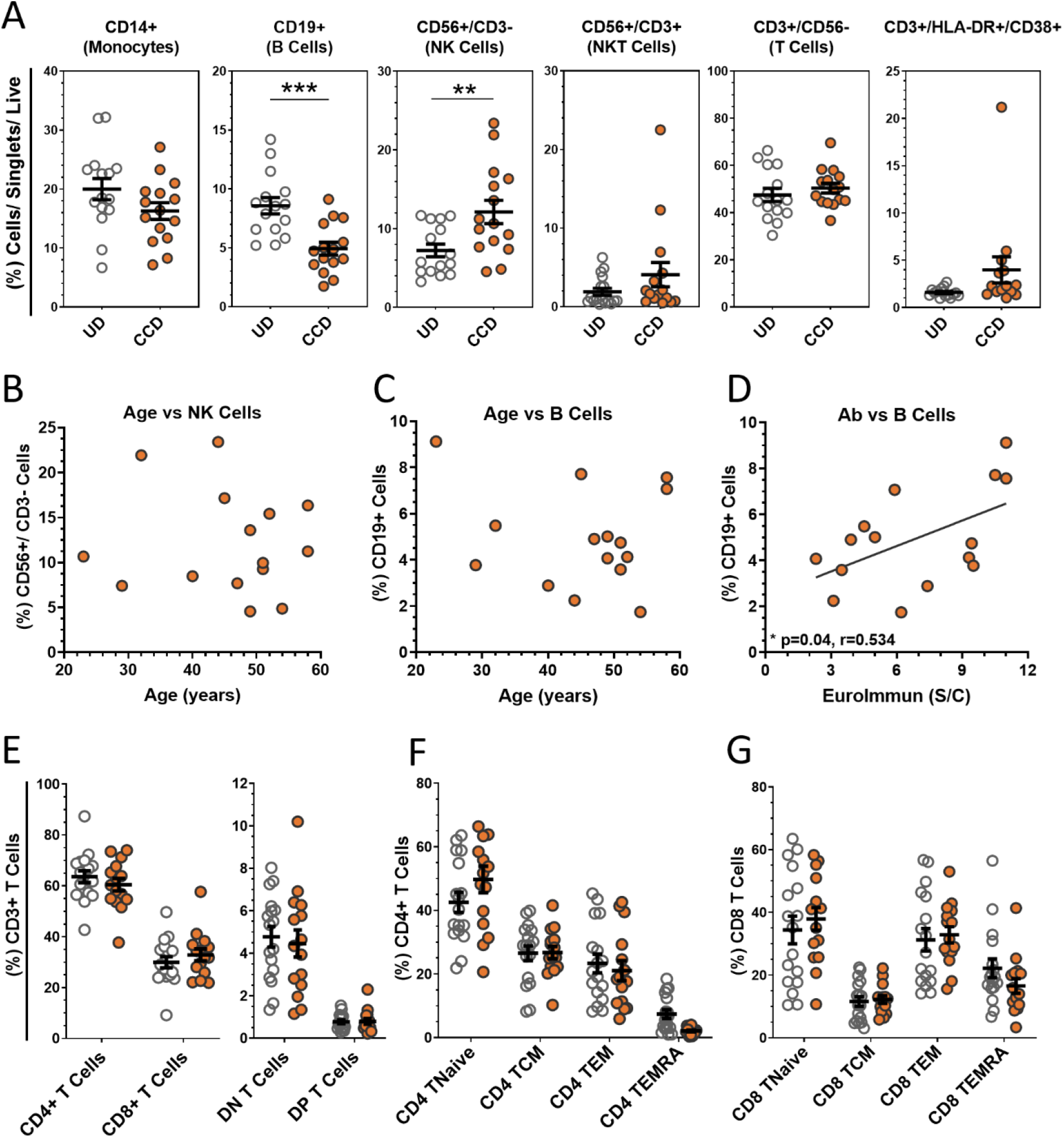
Analysis of COVID-19 convalescent donor buffy coat-derived PBMCs. **(A)** Buffy coat-derived PBMCs from COVID-19 convalescent donors (CCD, n=15, orange circles) and healthy uninfected donors (HD, n=17, clear circles) were assessed for leukocyte lineage by flow cytometry (see supplementary figure S1A for gating strategy). No correlation was seen between age of COVID-19 convalescent donors with **(B)** NK cells or **(C)** B cells, but significant correlation between **(D)** SARS-CoV-2 serum antibody content and the percentage of B cells between donors (p=0.04, r=0.534, Pearson correlation coefficient). **(E)** Analysis of the T cell compartment (see supplementary figure S1B for gating strategy) shows comparable mean levels of T cell subtypes, as well as **(F)** CD4+ and **(G)** CD8+ memory populations between HD and CCD. Data is represented as mean ± SEM. Significance determined by unpaired t-test with Holm-Sidak correction for multiple comparisons. *DN double negative (CD4−/CD8−), DP double positive (CD4+/CD8+), TNaive (CD62L+/CD45RA+/CD45RO−), TCM central memory (CD62L+/CD45RA−/CD45RO+), TEM effector memory (CD62L−/CD45RA−/CD45RO+), TEMRA terminal effector memory CD45RA revertant (CD62L−/CD45RA+/CD45RO−).*

**Table 1.** Baseline characteristics of COVID-19 convalescent donors and immune response at donation. Antibody level refers to Euroimmun assay values (>1.1 = positive). *Percentage CD3+/IFN-γ cells responding to combined SARS-CoV-2 pooled peptides (cf Figure 3).

| Donor Code | Blood Group | Days from symptoms onset to resolution | Days from symptoms resolution to donation | Antibody level | % SARS-CoV-2 VST* |
| --- | --- | --- | --- | --- | --- |
| C19BC1 | O pos | 8 | 38 | 10.5 | 0.22 |
| C19BC2 | O pos | 10 | 36 | 5.9 | 0.22 |
| C19BC3 | A pos | - | 34 | 9.3 | 0.13 |
| C19BC4 | O pos | 13 | 45 | 3.1 | 0.12 |
| C19BC5 | O pos | 14 | 52 | 3.9 | 0.03 |
| C19BC6 | A neg | 14 | 55 | 2.3 | 0.05 |
| C19BC7 | O pos | 14 | 56 | 6.2 | 0.07 |
| C19BC8 | B pos | 10 | 46 | 9.5 | 0.10 |
| C19BC9 | O neg | 18 | 37 | 7.4 | 0.16 |
| C19BC10 | O pos | - | - | 11 | 0.26 |
| C19BC11 | B neg | 7 | 49 | 11 | 0.05 |
| C19BC12 | A pos | 7 | 53 | 9.42 | 0.13 |
| C19BC13 | O pos | - | - | 5 | 0.08 |
| C19BC14 | O pos | - | - | 4.51 | 0.07 |
| C19BC15 | O neg | 13 | 53 | 3.48 | 0.03 |

### CCD T cell responses to spike, nucleocapsid and membrane SARS-CoV-2 peptides

PBMC were stimulated with SARS-CoV-2 peptide pools for spike protein, nucleocapsid protein and membrane glycoprotein or combined pools of all three and subsequently labelled for T cell surface markers (CD3, CD4, CD8) and intracellular cytokines (IFN-γ, TNF-α, IL-2) or activation markers (CD38, CD154, CD137). Representative flow analysis for a UD and CCD stimulated with combined SARS-CoV-2 peptide pools, with gating applied from a no-antigen control, is shown in Figure 2A. The percentage of SARS-CoV-2 VSTs in CCD positively correlated (Pearson correlation p=0.0381, r=0.5391) with SARS-CoV-2 antibody level (Fig 2B). Interestingly, the percentage of SARS-CoV-2 VSTs in CCD was found to decline significantly over time (Pearson correlation p=0.0021, r=0.6275) (Fig 2C). The mean percentage of CD3+ cells expressing IFN-γ, TNF-α and CD154 (Fig 2D) was significantly higher in CCD compared to UD for stimulation with each individual peptide pool and also for the combined peptide pools. CCD T cell IFN-γ, TNF-α and CD154 responses to individual peptide pools (n=10) was compared using repeated measures (RM) one-way ANOVA to determine whether there was a preferential response to specific SARS-CoV-2 antigens. While the mean percentage of CD3+/IFN-γ+ cells and CD3+/TNF-α+ cells was comparable between the three peptide pools, the CD4+/CD154+ response was significantly higher (p=0.042) to membrane peptides than to nucleocapsid peptides. Altogether, these data indicate there is no consistent preferential T cell cytokine response to one particular SARS-CoV-2 antigen (Fig S3A).

**Figure 2.**
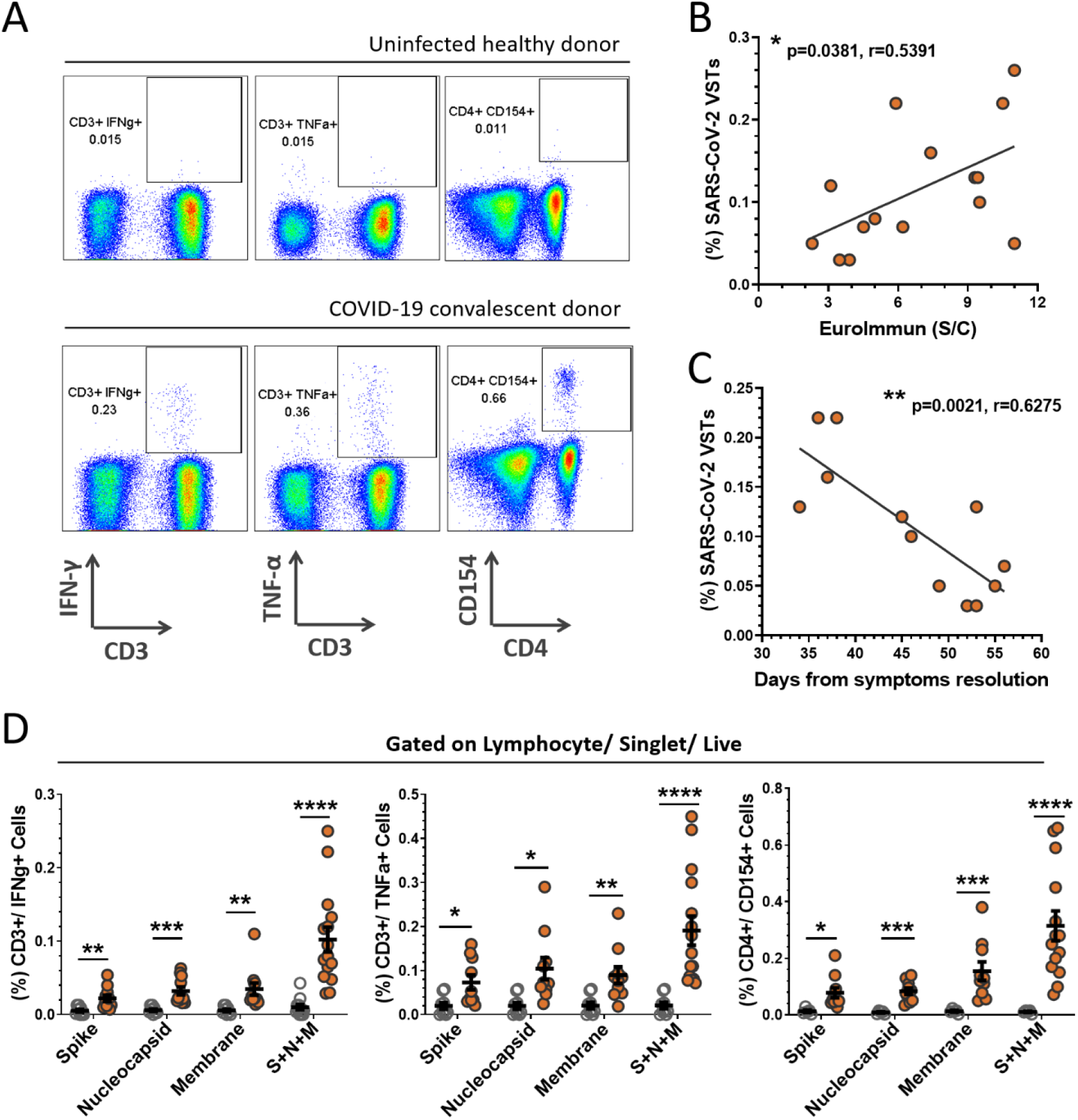
PBMC responses to SARS-CoV-2 peptides. PBMCs derived from buffy coats were incubated with SARS-CoV-2 peptides (Spike + Nucleocapsid + Membrane) for 5 hours and corrected against a no antigen control well for positive expression of cytokines and activation markers. **(A)** Representation of flow cytometric analysis from a healthy uninfected donor (HD) and COVID-19 convalescent donor (CCD), note all flow analyses were gated on lymphocytes/ single cells/ live cells and subsequently quantified for percentage CD3+/IFN-γ+ cells, CD3+/TNF-α+ cells and CD4+/CD154+ cells (S2 for gating strategy). The percentage of SARS-CoV-2 VSTs in the CCD PBMC population (i.e. CD3+/IFN-γ+ cells reactive to pooled S+N+M peptides corrected to no antigen control) significantly correlated with **(B)** antibody titre at donation (p=0.0381, r=0.5391) and **(C)** days from resolution of symptoms to donation (p=0.0021, r=0.6275). Calculation was performed using Pearson correlation coefficient. **(D)** Mean percentages of CD3+/IFN-γ+ cells, CD3+/TNF-α+ cells and CD4+/CD154+ cells for individual and pooled peptides corrected to no antigen control were compared between HD (n=12, clear circles) and CCD (n=15, orange circles). Data is represented as mean ± SEM. Statistical significance was determined using unpaired t-tests corrected for multiple comparisons using the Holm-Sidak method where *p≤0.05, **p≤0.01, p≤0.001 and **** p≤ 0.0001.

Further dissection of the cytokine response to SARS-CoV-2 peptide pools within lymphocyte subsets CD4+ T cells, CD8+ T cells and NK cells (CD56+/CD3− PBMCs) from CCD indicates the IFN-γ response is primarily by CD4+ T cells (Fig S3B). The mean percentage of CD4+/IFN-γ+ PBMC was significantly higher than either CD8+/IFN-γ+ or CD56+/IFN-γ+ cells for each individual peptide pool. Stimulation with combined peptide pools drove induction of higher percentages of CD4+/IFN-γ+ T cells than either CD8+/IFN-γ+ or CD56+/IFN-γ+ cells. The percentage of CD8+/IFN-γ+ cells was significantly increased over CD56+/IFN-γ+ cells. Conversely the TNF-α response to pooled peptides demonstrated significantly higher CD56+/ TNF-α+ cells than CD8+/ TNF-α+ cells (Fig S3C).

### Isolation of SARS-CoV-2 VSTs using peptide-driven IFN-γ selection and expansion in culture

PBMC were stimulated with combined peptide pools and reactive VSTs were isolated with the CliniMACS IFN-γ Cytokine Capture System (CCS) kit. Analysis of the IFN-γ selected T cells (Figure 3A) revealed an equal ratio of monocytes to T cells with negligible levels of NK or NKT cells. CD3+ T cells in the isolated fraction were a mix of CD4+ (53.02 ± 3.94%) and CD8+ (35.73 ± 3.23%) cells; where CD4+ T cells were predominantly central memory (86.52 ± 3.44%), CD8+ T cells showed mostly effector memory and terminal effector RA (TEMRA) phenotype. The non-target cells from the CCS isolation were irradiated and co-cultured with the isolated IFN-γ+ target cells to act as feeders for VST culture expansion in G-Rex culture vessels. After 14 days expansion (Fig 3B), cultures were highly enriched for T cells (87.95 ± 2.99%) with minimal expansion of NK and NKT cells. T cells were predominantly CD4+ (77.86 ± 5.19%) with smaller proportion of CD8+ T cells (18.05 ± 4.4%). Both the CD4+ and CD8+ populations were heavily skewed towards central memory phenotype. Direct comparison of populations between isolation and day 14 expansion showed significant differences in monocyte and T cell content, CD4+, CD8+ and Double Negative (DN) T cells, and memory subpopulations in both the CD4 and CD8 compartment (Fig S4) demonstrating an enrichment of central memory CD4 cells in our culture process.

**Figure 3.**
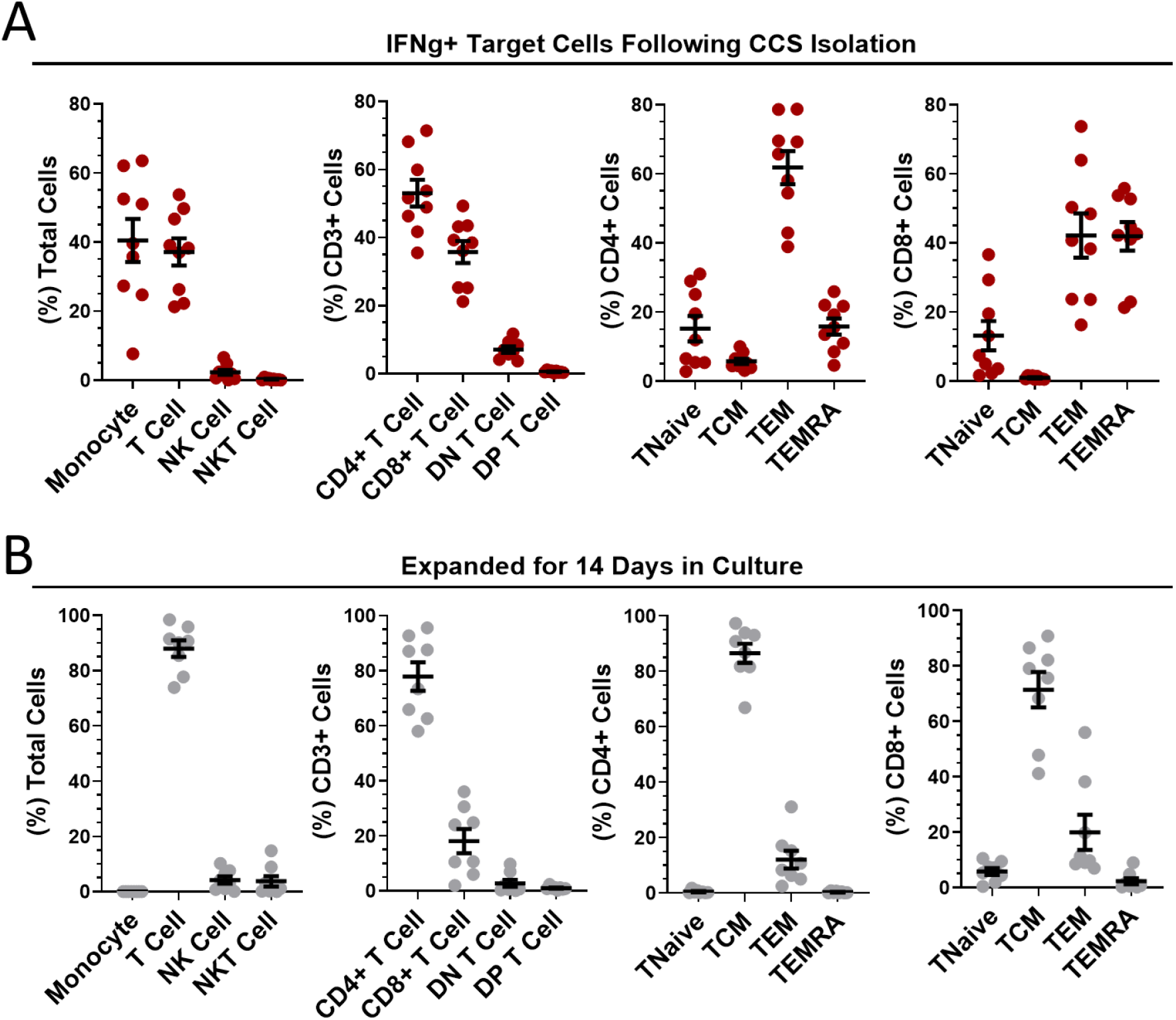
Phenotypic analysis of isolated and expanded SARS-CoV-2 VSTs. The percentages of leukocytes, T cell subpopulations and CD4/CD8 differentiation status were quantified for **(A)** IFNg+ target cells directly after SARS-CoV-2 peptide-mediated cytokine capture system (CCS) isolation and **(B)** following expansion in culture for 14 days. All data is represented as mean ± SEM.

### Expanded SARS-CoV-2 VSTs show specific response to all SARS-CoV-2 peptides

SARS-CoV-2 peptide pool-loaded dendritic cells (DCs) and unloaded DC controls were then co-cultured with 14-day expanded VST at [1 DC: 10 VST] and analyzed for T cell activation and cytokine expression. Both CD4+ and CD8+ VSTs demonstrated specific anti-viral reactivity via expression of IFN-γ, TNF-α, CD154, CD107a and CD137 when co-cultured with autologous DC loaded with SARS-CoV-2 pooled peptide (representative plots Fig 4A). There was a stronger response to peptide re-stimulation in CD4+ T cells than in the CD8+ T cells for IFN-γ, TNF-α, CD154 (Fig 4B), but equivalent CD107a and CD137 expression. The total T cell response to? IFN-γ and TNF-α response to individual pools for each donor VST are shown in Fig 4C and Fig 4D respectively, indicating donor-specific variation. When the data was collated, equivalent reactivity to all three peptide pools was observed (Fig 4E). Although the total T cell population did not show predominance for any of the SARS-CoV-2 antigen pools, the IFN-γ response was also assessed for the individual peptide pools, gated on CD8+ T cells and CD4+ T cells specifically (representative plot Fig 4F). In CD4+ T cells there was a similar response to each peptide pool, but in CD8+ T cells, the nucleocapsid peptide pool was clearly immunodominant, inducing the strongest response (Fig 4G and H).

**Figure 4.**
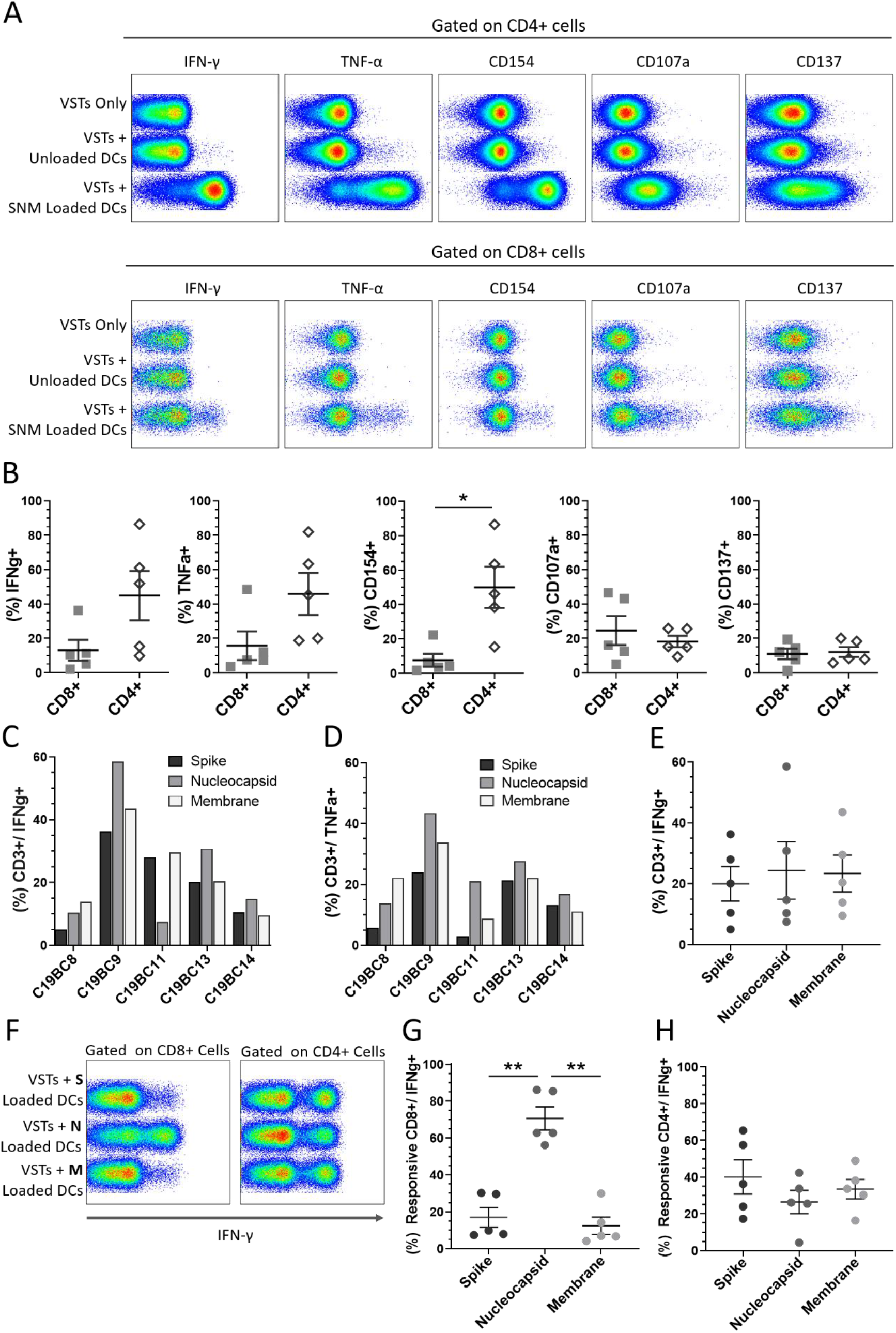
Cultured SARS-CoV-2 VST peptide specificity. Isolated and expanded SARS-CoV-2 VST (n=5) at day 14 culture were co-cultured with peptide-loaded mature autologous DC. **(A)** Flow cytometric analysis on either CD3/CD4+ or CD3/CD8+ cells for IFN-γ, TNF-α, CD154, CD107a and CD137 is shown for negative controls (VSTs Only, and VSTs + Unloaded DCs), and VST with pooled SARS-CoV-2 peptide loaded DCs (VST + SNM-loaded DCs). **(B)** The mean percentage of CD4+ and CD8+ positive for each marker of DC-stimulated VST was compared using paired t-test Holm-Sidak correction for multiple comparisons. *p≤0.05. **(C, D)** Individual donor VST were assessed for T cell response (% CD3+/IFN-γ+ and % CD3+/TNF-α+ respectively) against DCs loaded with individual SARS-CoV-2 peptide pools: spike, nucleocapsid and membrane. **(E)** Collated responses to the individual peptides in the total CD3+ population indicated no significant difference. **(F, G)** A significantly higher CD8+/IFN-γ+ cells response was seen with nucleocapsid stimulation than with the other peptide pools (significance determined using RM one-way ANOVA with Geisser-Greenhouse correction **p≤0.01). **(H)** CD4+/ IFN-γ+ cells responded similarly to all the three peptide pools. All data represented as mean ± SEM.

We assessed whether other non-SARS virus-specific T cells were coincidentally expanded in this process. Stimulation with EBV, adenovirus or irrelevant (GAD65) peptides did not demonstrate any significant levels of T cells directed to other viruses as measured by intracellular IFN-g response (Fig S5).

### Culture Optimization to Enhance SARS-CoV-2 VST Expansion for Clinical Manufacture

Adoptive transfer of VSTs relies upon significant cell expansion in order to provide sufficient doses for clinical trial or therapeutic treatment. Combined growth curves of VST samples C19BC8-C19BC14 (n=6) cultured at optimized seeding density demonstrated a 2-3 log expansion from the initial isolated IFN-γ+ cells at day 0 to day 14, followed by a general plateau in expansion beyond this point (Fig 5A). Additionally, some VST cultures were expanded in different medium supplements for optimization of culture conditions, supplemented with IL-2, IL-7 or commercial pathogen-inactivated human platelet lysate (hPL). Addition of IL-7 had no effect on culture expansion between day 0 and day 8 (Fig 5B), whereas addition of hPL induced a markedly higher fold expansion between day 0 to day 14 compared to IL-2 alone in each donor culture tested (Fig 5C). After day 14 culture expansion plateaued, and an increased transition from central memory to effector memory phenotype by day 21 was observed (representative plot Figure 5D). When cultures were administered a second feeder cell re-stimulation (FR) with autologous irradiated cells at day 14, central memory phenotype was retained at day 21. FR induced a subsequent two log expansion between day 14-21 (Fig 5E) in all VST cultures tested (n=5). The final VST numbers harvested under optimized conditions from a single CCD buffy coat ranged from 1-4.6×10^9^ at day 14, and 0.3-2×10^11^ at day 21 following FR (Fig 5F). Cultures were monitored throughout expansion to determine whether FR affected culture composition, but no significant differences in lymphocyte subsets (Fig 5G-H) or T cell memory status (Fig 5I-J) were observed between cells harvested at day 14 or at day 21.

**Figure 5.**
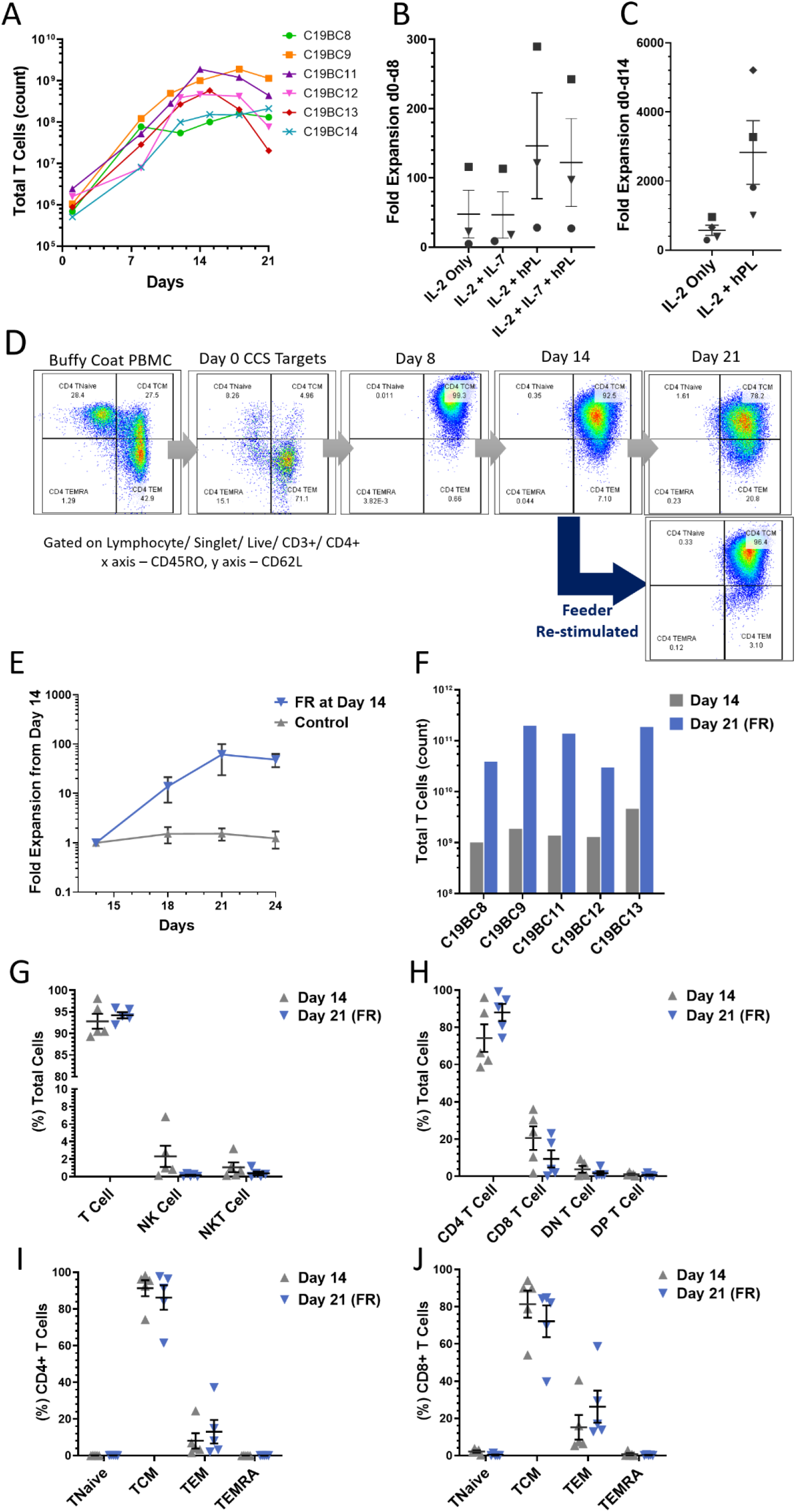
SARS-CoV-2 VST culture optimization. **(A)** Isolated SARS-CoV-2 VST from donors C19BC8-14 had a 2-3 log expansion over 21 day culture using an optimized culture expansion protocol. Variation in the start numbers of VST reflect donor variation in initial buffy coat PBMC numbers. Fold expansion between **(B)** day 0 and day 8 and **(C)** day 0 and day 14 was assessed in cultures after supplementation with IL-2, IL-7 and human platelet lysate (PL). Donors C19B9 (square), C19BC11 (triangle), C19BC12 (circle) and C19BC13 (diamond) were divided to compare medium supplementation condition. (**D**) Representative culture C19BC9 by day 21 without re-stimulation indicated some transition of CD4 TCM to CD4 TEM (Day 21 top panel). CD4 CM phenotype was retained when cultures re-stimulated at day 14 with autologous irradiated feeders (Day 21 bottom panel). **(E)** VST cultures were split at day 14 to directly compare standard continuation in culture (control) and re-stimulation with autologous irradiated feeders. Data is represented as mean T cell count ± SEM (n=5). **(F)** VST from a single donor buffy coat were compared for optimal cell yields at day 14 (grey), and day 21 with feeder re-stimulation at day 14 (Day 21 FR, blue). **(G-J)** Final product phenotype and T cell memory status was compared at both harvest time-points: Day 14 (grey circles) and Day 21 FR (blue triangles). Data is represented as mean ± SEM. No significant differences were observed using paired t-tests with Holm-Sidak correction for multiple comparisons.

## Discussion

The recent global outbreak of SARS-CoV-2 infection has caused significant issues in many countries due to the strains on both healthcare infrastructure and economic climate. This viral infection is associated with widely divergent degrees of clinical severity, with most infected people having asymptomatic or mild symptoms, but severe effects in a proportion of individuals. These infections have been extensively studied and reported and have indicated that failure of key aspects of the immune response appear to be consistent with progression to severe disease and mortality. The principal features include alterations to the neutrophil: lymphocyte ratio (11, 23), loss of T cells (24), and increased expression of exhaustion and senescence markers within the T cell compartment (10, 25, 26).

Adoptive anti-viral T cell therapy has been an important therapeutic approach for other infections such as EBV, CMV and adenovirus (20, 22, 27). Various manufacturing methods have been developed, but recently cytokine capture has proved effective for isolation of T cells for clinical therapy (17–19).

PBMC fractions from whole blood (buffy coats) from CCD confirmed by SARS-CoV-2 RNA assay (n=15) were collected between 34-56 days after resolution of symptoms, and only patients with mild symptoms (non-hospitalized) were used, as current reports suggest that in these patients a robust T cell response is generated and maintained (28), without the aberrant T cell phenotype seen in severe infections. The immunophenotyping of PBMC demonstrated broadly similar percentages of different immune subpopulations compared to UD, though there was a significant decrease in CD19 B cells in CCD in common with many other reports (11, 25). However, there was a significant increase in the NK compartment in CCD. Large cohort studies investigating immunosenescence of healthy individuals report a concomitant decline in T and B cells with preservation or increase in NK cell numbers (29), indicating a more important role for these innate immune cells to clear infections in elderly people. Though increased innate lymphocyte levels have been correlated with increasing age, there was no correlation between age and frequency of NK cells or B cells. There was a significant correlation between B cell level and SARS-CoV-2-specific antibody levels. The T cell compartment showed no significant differences in CD4/8 ratios or differentiation status between CCD and UD, which indicates that mild COVID-19 does not significantly affect the overall T cell composition There is however clear evidence of double-positive T cells, associated with viral infections (30).

The T cell response in CCD stimulated with spike, nucleocapsid and membrane glycoprotein peptide pools demonstrated that there was a small but distinct population of virus-specific T cells present, which were absent in UD. A number of reports have indicated that protein or peptides from the C terminal of the spike protein can elicit T cell responses in donors known to be SARS-CoV-2 negative, indicating a cross-reaction with conserved motifs in other coronaviruses (31, 32) but this was not observed using this spike peptide pool. There was some indication that peptide pools from different proteins elicited a differential cytokine response, with IFN-γ and TNF-α responses stronger to nucleocapsid peptides, though CD154 was preferentially increased in response to membrane peptides. This correlates well with findings in other cases of mild COVID-19 (33). The relatively low percentage of VSTs detectable has been reported in other studies on COVID-19 patients with mild disease (31), but there is evidence that SARS-CoV-2 infection blunts the cytokine response in a wide variety of leukocytes (34). The key finding from this was that we could successfully elicit IFN-γ responses in SARS-CoV-2 VSTs from CCD peripheral blood, which then determined that we could isolate and expand these T cells using an established clinical-grade cytokine capture assay. The principal confounding factor was identifying that SARS-CoV—2 VST levels were closely correlated with time from resolution of infection, with VSTs dropping to less than 0.03% by 60 days post-resolution of infection. Further work is ongoing to determine whether SARS-CoV-2-VST remain detectable later in convalescence, though subsequent re-exposure to the virus may result in reinforcement and expansion of residual cells.

Leucocytes isolated using Good Manufacturing Practice (GMP)-compliant IFN-γ bead selection resulted in a mixed cell population with monocytes and NK cells as well as the SARS-CoV-2 VSTs, but after culture in closed-process flasks, there was a rapid expansion of the T cells, and a significant modulation of T cell phenotype. The initial peptide-reactive T cells were predominantly differentiated effector T cells, with an equal CD4:CD8 split, but after 14 day culture, there was an overwhelming shift to central memory phenotype with a strong skew to CD4 T cells. This change may reflect a loss of effector T cells and a massive expansion of the central memory (TCM) compartment. There is retention of few NK cells, though the monocytes are lost during culture. The cultured VSTs also retain strong specificity for viral peptides, as shown by the functional assay of response, where co-culture with autologous DC drives a pronounced CD4 activation and cytokine response, indicating that these expanded T cells would have a significant supportive role in recipients. Interestingly, the CD8 response was significantly lower than CD4 VSTs, with low expansion and weaker responses to the SARS-CoV-2 antigens. This lower response may be an advantage for a therapeutic product, as in models there is a clear protective role for CD8 T cells against acute SARS infection (35), but there is evidence that hyper-activation of CD8 T cells is linked to severity of COVID-19 disease (36). The expanded CD8 VST cell demonstrated differential responses to each protein peptide pool and there was clear indication that the nucleocapsid protein is the immunodominant antigen for the cytotoxic T cell population.

The initial 14 day culture of SARS-CoV-2 VSTs in GMP-compliant reagents demonstrated that a suitable therapeutic T cell product could be manufactured from even small numbers of VSTs present in a single unit of blood, yielding up to 5×10^9^ cells per manufacturing run with greater than 90% central memory T cells and a 3.3 log expansion. However, in common with other VST cells cultures, growth peaked at day 14-15 and tended to decrease by day 21, with concomitant changes in the differentiation status of the T cells. Supplementation of the medium with IL-7 had no significant effect on growth or phenotype, but addition of hPL significantly increased T cell expansion, suggesting that other cytokines and growth factors may support and drive T cell proliferation. Further expansion was provided by a second round of stimulation with autologous feeder cells, resulting in an approximate 2-log further increase in cell numbers with a consolidation of central memory phenotype, plus reduction in NK cells. There was a further skew towards CD4 T cells during this second expansion. Thus, with the use of leukapheresis instead of buffy coats, fully-closed isolation and expansion using CliniMACS Prodigy cell processing and G-Rex culture flasks we could generate sufficient material to treat multiple patients from a single suitably HLA-matched donor, with or without a second expansion phase. HLA matching is important for effective function of adoptive T cell therapies, and it is clear that some HLA types have increased susceptibility to COVID-19 (37), so supplementation of immune response with donor T cells from a matched donor could be a therapeutic option where few others exist.

It is now clear that there are several immunotypes of patients with COVID-19, some of which are predicated towards more severe infection and higher mortality (38). The GMP-compliant T cell isolation and expansion process we have outlined clearly demonstrates that CCD with mild symptoms do not have the disruption to the T cell compartment seen in severe cases. VSTs can be isolated and rapidly expanded to sufficient yield to provide multiple doses for adoptive T cell therapy in patients with symptoms severe enough to require hospitalization. Therefore, it is feasible to generate T cell products for clinical trials in support of severe SARS-CoV-2 infections where the endogenous T cell response is compromised, representing a potentially significant advance in therapy for COVID-19.

### Limitations of this study

This study has used the 3 SARS-CoV-2 Ag peptide pools available from April 2020 to generate the phenotyping and expansion data of antigen-specific T cells. Additional antigens may reveal a fuller picture of the immune response in COVID-19, although their addition to the T cell expansion method described here could have a positive additive effect. The demographics of the local donor pool used is necessarily limited to blood donors who have undergone mild COVID-19 disease. Different T cell responses and proliferation characteristics may be found in other donor populations recovering from a range of COVID-19 symptoms. Clinical use of this product will require controlled clinical trials for safety and efficacy assessment.

## MATERIALS & METHODS

### Study Design

The aim of this study was to investigate the potential to isolate and expand SARS-CoV-2 peptide-specific T cells from CCD and evaluate this for use as an allogeneic ‘off the shelf’ VST therapy for hospitalized COVID-19 patients. To explore this, CCD were recruited through the SNBTS Convalescent Plasma program. CCD were eligible if they had a confirmed positive SARS-CoV-2 PCR test and were a minimum of 28 days after resolution of infection symptoms, as well as fulfilling the current criteria for whole blood donation. UD (adults confirmed as having suffered no COVID-19 symptoms at time of donation) were used to compare initial phenotyping and SARS-CoV-2 antigen T cell responses with CCD. Buffy coats from CCD (n=15) and UD (n=17) were obtained under Sample Governances 20~02 and 19~11 respectively. All donations were fully consented for research use.

#### SARS-Cov-2 antibody detection

The Euroimmun anti-SARS-CoV-2 assay (Euroimmun US, NJ, USA) clinical diagnostic indirect ELISA was used to detect antibodies to SARS-CoV-2 spike protein from donor serum according to the manufacturer’s instructions. The results were expressed as a ratio against a calibrator control, where values <0.8 is considered negative and >1.1 are considered positive.

### Buffy Coat Peripheral Blood Mononuclear Cell Isolation

Buffy coats were diluted [1:3] with PBS and added to Leukosep tubes containing Ficoll-Paque (GE Healthcare). Tubes were centrifuged at 450g for 40 minutes and the resulting buffy layer extracted. Isolated PBMCs were then washed in PBS and counted on MACSQuant10 Analyzer (Miltenyi Biotec).

### Immunophenotyping

PBMC directly after isolation from buffy coats, and T cells from VST cultures (see below) were analyzed for surface immunophenotype. For this, 2×10^6^ cell samples were taken and washed with PBS buffer supplemented with EDTA and human serum albumin (PEA buffer). Cell pellets were re-suspended in 100 μL PEA and incubated with 5μL Fc Receptor blocking reagent to prevent non-specific antibody binding. Antibody surface marker multi-color panels detailed in Fig S1 were then added for 20 minutes at 4°C. Samples were then washed and re-suspended in PEA, with dead cell dye DRAQ7 (eBioscience) added prior to acquisition on a MACSQuant10 Analyzer (Miltenyi Biotech) recording a minimum of 100,000 events.

### PBMC Stimulation with SARS-CoV-2 Peptides

SARS-CoV-2 Peptivator peptide pools (Miltenyi Biotech) containing 15-mer sequences with 11 amino acids overlap for the immunodominant section of the spike protein, and the full sequence for nucleocapsid protein and membrane protein were reconstituted in DMSO/water according to manufacturers’ guidelines.

PBMC were plated in TexMACS medium (Miltenyi Biotech) in a 24-well plate at 5×10^6^ cells/mL per well with treatments: negative control, PMA/ionomycin positive control, individual spike, nucleocapsid, and membrane peptide pools, and combined pools (spike + nucleocapsid + membrane - SNM). SARS-CoV-2 peptide pools were used at [0.3 nmol/mL], and cell activation cocktail (BioLegend) added to the positive control well at [1X]. The negative control well contained DMSO/water at the same volume as the peptide wells. Cells were stimulated for a total of 5 hours at 37°C, 5% CO2; with Brefeldin A (BioLegend) added at [5 μg/mL] for the final 3 hours.

### Intracellular Labelling

Plates were harvested into FACS tubes and washed with PEA. Samples were treated with Fc Receptor blocking reagent as above, and surface marker antibodies for multi-color panels (see Figure S2 for details) were then added for 20 minutes at 4°C. Cells were then washed and stained with fixable viability dye (FVD) eFluor780 (eBioscience) for 30 minutes at 4°C. Cells were subsequently fixed and permeabilized using Cytofix/perm kit (BD Biosciences) for 20 minutes at 4°C, then washed and labelled with antibodies for intracellular cytokines (detailed in Supplementary table 2) for 20 minutes at 4°C. Cells were washed and analyzed with a MACSQuant10 Analyzer recording a minimum of 150,000 events.

### VST Isolation

SARS-CoV-2 VSTs were isolated from PBMC whole population using a cytokine capture system (CCS) assay. Briefly, PBMC were plated at 5×10^6^ cells/mL/cm^2^ in standard corning plates and incubated overnight at 37°C, 5% CO_2_. The following morning, plates were stimulated with pooled SARS-CoV-2 peptide pools (spike + nucleocapsid + membrane) each at [0.3nmol/mL] for 6 hours at 37°C, 5% CO_2_. The virus-specific IFN-γ secreting cells were then isolated using the IFN-γ CCS assay by either manual column or CliniMACS Prodigy isolation as described in (17). Following isolation, each fraction was counted and phenotyped using the lymphocyte panel as described above and illustrated in Fig.S1.

### VST Culture Optimization

Non-target cells from the IFN-γ CCS assay were irradiated at 40Gy and used as feeders for the IFN-γ+ target cells. Cultures were initially seeded at either 1×10^7^ total cells per cm^2^ [200-400 non targets: 1 target], or 3×10^6^ total cells per cm^2^ [100 non targets: 1 target] in G-Rex culture vessels (Wilson Wolf). Cells were cultured in GMP-grade TexMACS medium, and supplemented to determine culture optima using [200U/mL] IL-2 (GE Healthcare), [155U/mL] IL-7 (Miltenyi Biotech), [2%] human AB Serum (SNBTS) or [2%] nLiven (Cook Regentec). Cells were cultured for up to 28 days with feeds every 3-4 days and cultures split as necessary to maintain a density of 0.5-3×10^6^ T cells/cm^2^. At day 14, VST cultures from six donors were split to test feeder re-stimulation, where thawed irradiated non-target cells were added to cultured VSTs at [10 non targets: 1 VST] alongside a control culture with no feeder re-stimulation. Samples were taken every 3-4 days for immunophenotyping with the lymphocyte panel described above.

### Generation of Monocyte-Derived DC

Monocyte-derived DC were generated from isolated PMBC CD14+ monocytes. Briefly, monocytes were isolated from purified PMBC using anti-CD14 microbeads (Miltenyi Biotech) as per manufacturer’s instructions. Cells were cultured at 37°C, 5% CO2 for 6 days in RPMI (Life Technologies) supplemented with [5%] AB serum, [2mM] Glutamax (Sigma-Aldrich), [20 ng/mL] GM-CSF and [15 ng/mL] IL-4 (both Miltenyi Biotech). Media was replaced on days 2 and 4 of culture. After 6 days, cells were harvested using [1X] TrypLE (Life Technologies) and frozen in CryoStor 10 (Stem Cell Technologies) until required for VST stimulation.

### VST Stimulation Assay

Cells from VST cultures were taken at day 14 to test in a stimulation assay with peptide-loaded DC. Briefly, frozen autologous immature DC were thawed, plated with RPMI medium and stimulated with individual SARS-CoV-2 peptide pools and combined pools at [0.3nmol/mL] for 6 hours at 37°C, 5% CO_2_. Where possible, VSTs were tested for specificity to SARS-CoV-2 by testing against other virus-specific peptides (Adenovirus Hexon peptide, Epstein-Barr Virus consensus peptide pool, and GAD65 peptide, all Miltenyi Biotech). Unloaded DC were included as a negative control. DC were then washed and re-plated in RPMI supplemented with poly I:C [20 μg/mL] and PGE_2_ [1 μg/mL] at 2.5×10^5^ cells/cm^2^ overnight to drive DC maturation. The next morning DCs and VSTs were co-cultured at 2.5×10^6^/cm^2^ [10 VST: 1 DC]. A VST-only control well was used to measure baseline stimulation of VSTs co-cultured with unloaded DCs. Plates were then stimulated for 5 hours and labelled for the cytokine and activation panels as described above.

### Flow Cytometry Data Analysis

Analysis of all flow cytometry data was performed using either MACSQuantify (Miltenyi Biotec) for cell counts, or FlowJo version 7 (TreeStar Inc) for wider phenotyping analysis. All analyses were subject to a basic initial gating strategy in which debris was first excluded on the basis of forward and scatter properties, and sequentially gated for singlets using FSC Area versus Height, and finally sequentially gated on live cells (DRAQ7 or FVD negative cells). Populations and acquisition of activation markers/ cytokines were then quantified using percentages corrected to negative controls (see S1 and S2 for full gating strategies).

### Statistical Analysis

Statistical analysis was performed using GraphPad Prism version 8.4.2 (GraphPad Software). Comparisons of population frequencies between healthy donors and COVID-19 convalescent donors were performed using unpaired two-tailed Student’s t tests with Holm-Sidak correction for multiple comparisons. Tests comparing population frequencies intra-donor between time-points (i.e. day 0 versus day 14, and day 14 versus day 21) for lymphocyte phenotype markers or comparing acquisition of activation markers within the CD8 versus CD4 population were paired two-tailed Student’s T t-tests, corrected for multiple comparisons using the Holm-Sidak method where relevant. Response comparisons between the three individual SARS-CoV-2 peptides (spike, nucleocapsid and membrane) were tested using repeated-measures one-way ANOVA with the Geisser-Greenhouse correction to assume equal variability of differences within VST cultures. Correlation between intra-donor frequency of populations compared to other baseline characteristics was done by computing linear correlation coefficients using Pearson’s correction with confidence intervals of 95%. Unless otherwise stated, data is represented as mean values ± SEM.

## Acknowledgments

The authors wish to express their gratitude to all blood donors who volunteered to provide their blood during this unpreceded health crisis. We thank the whole SNBTS donor, processing, testing, medical and nursing teams for their support. We thank the NMRU and H and I labs for advice and patient testing. We thank Dr Anne Richter of Miltenyi Biotec GmbH for scientific advice around the use of the peptide pools.

## Author Contributions

RSC, ARF, MLT and JDMC conceived and designed the study. RSC, ARF, MLT and JDMC drafted and revised the manuscript; RSC, ARF, LS, PB, SI performed experimental work. RSC, ARF, LS, PB, SI and JDMC analyzed data. LJ, SZ and MLT performed clinical tasks including identification, consenting and testing of donors. All authors contributed content and reviewed the manuscript.

## Data and materials availability

All data associated with this study are present in the paper or the supplementary materials.

## Supplementary Material

### Supplementary figures

**Figure S1.**
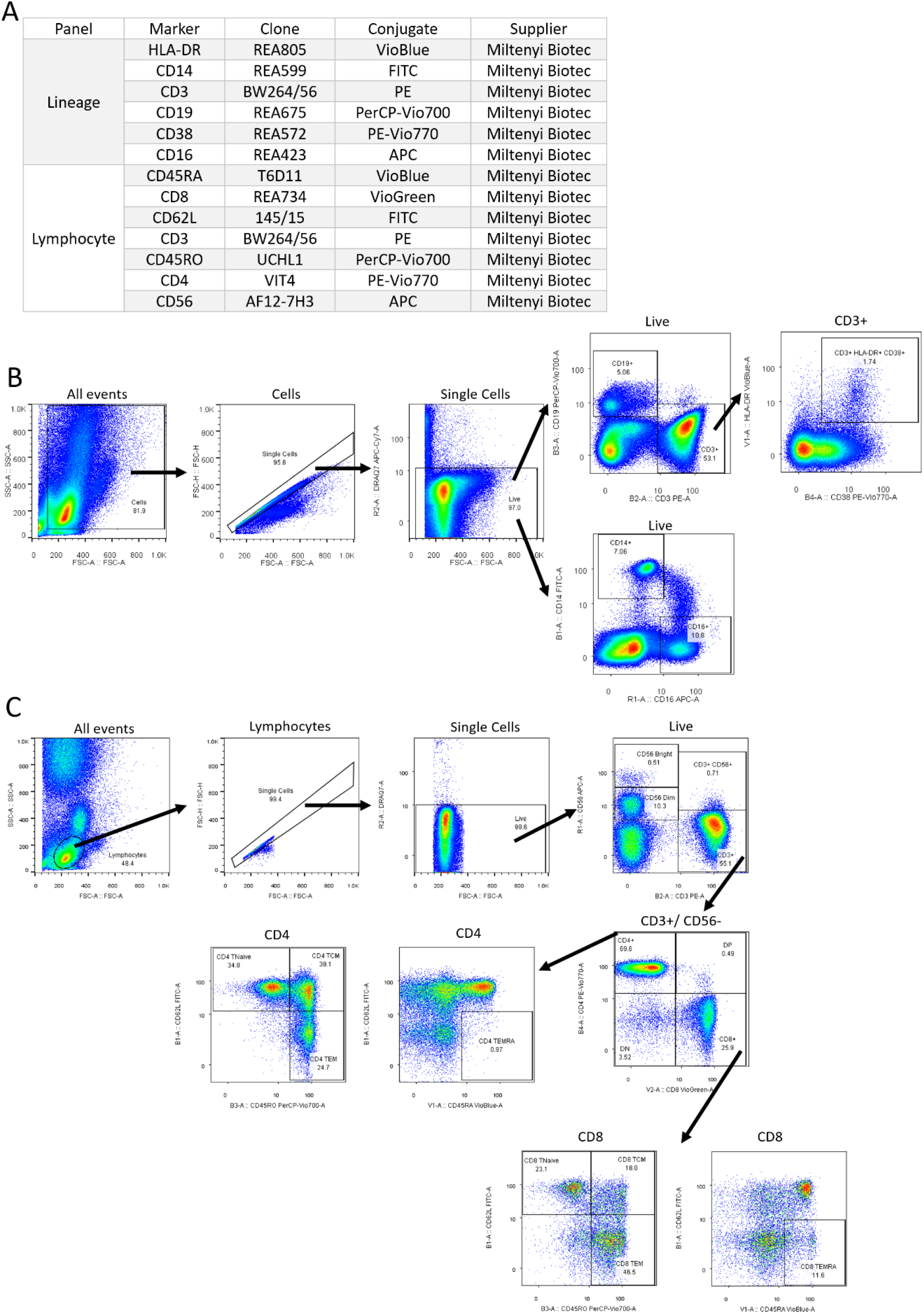
Surface phenotyping panels and gating strategies.

**Figure S2.**
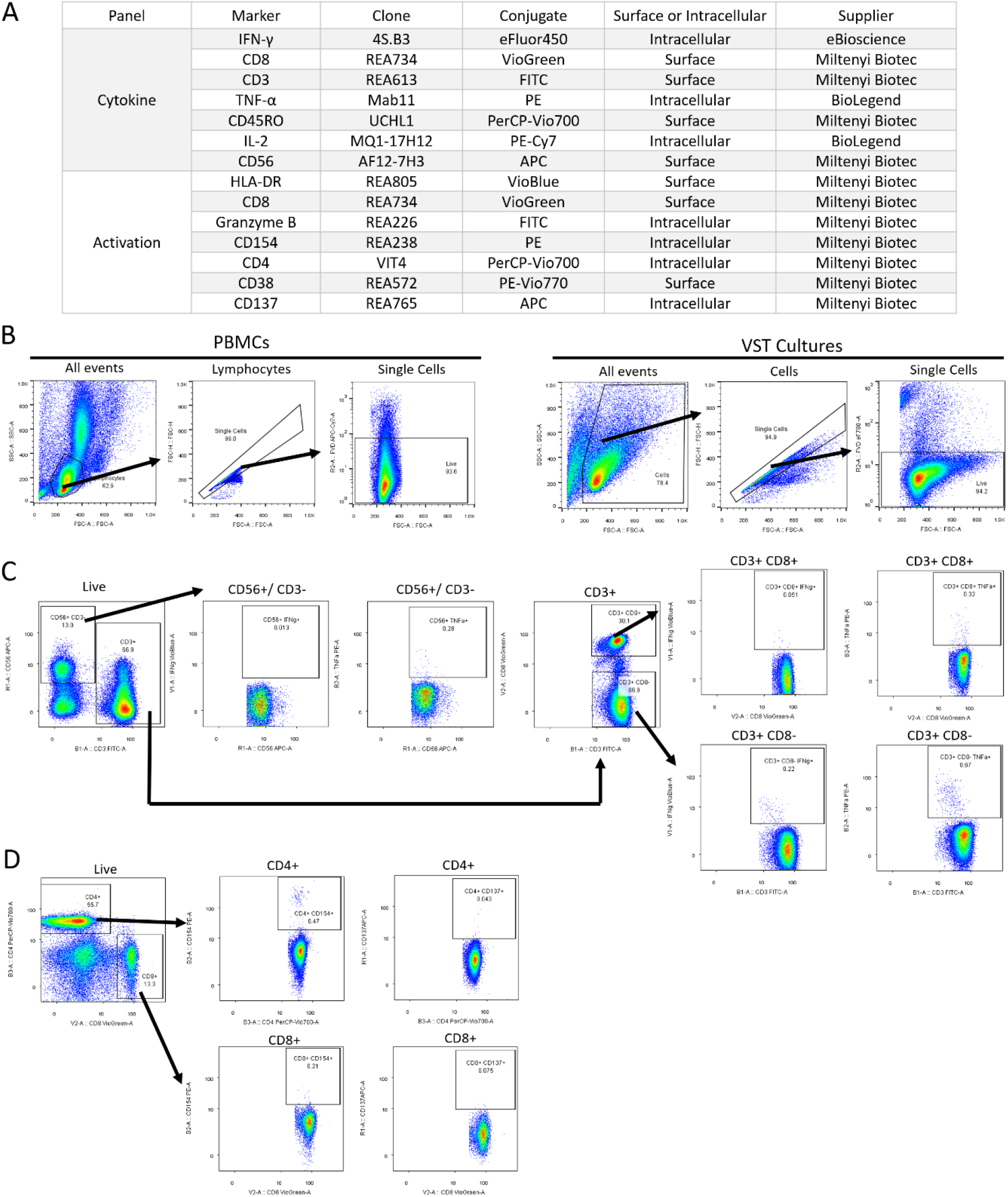
Intracellular flow cytometry panels and gating strategies. **(A)** Antibodies used in cytokine and activation panels. **(B)** All analyses were subject to initial sequential gating as shown for PBMCs and VST cultures. Flow cytometry gating strategies following initial gating above for **(C)** cytokine and **(D)** activation panel are shown using representative PBMC.

**Figure S3.**
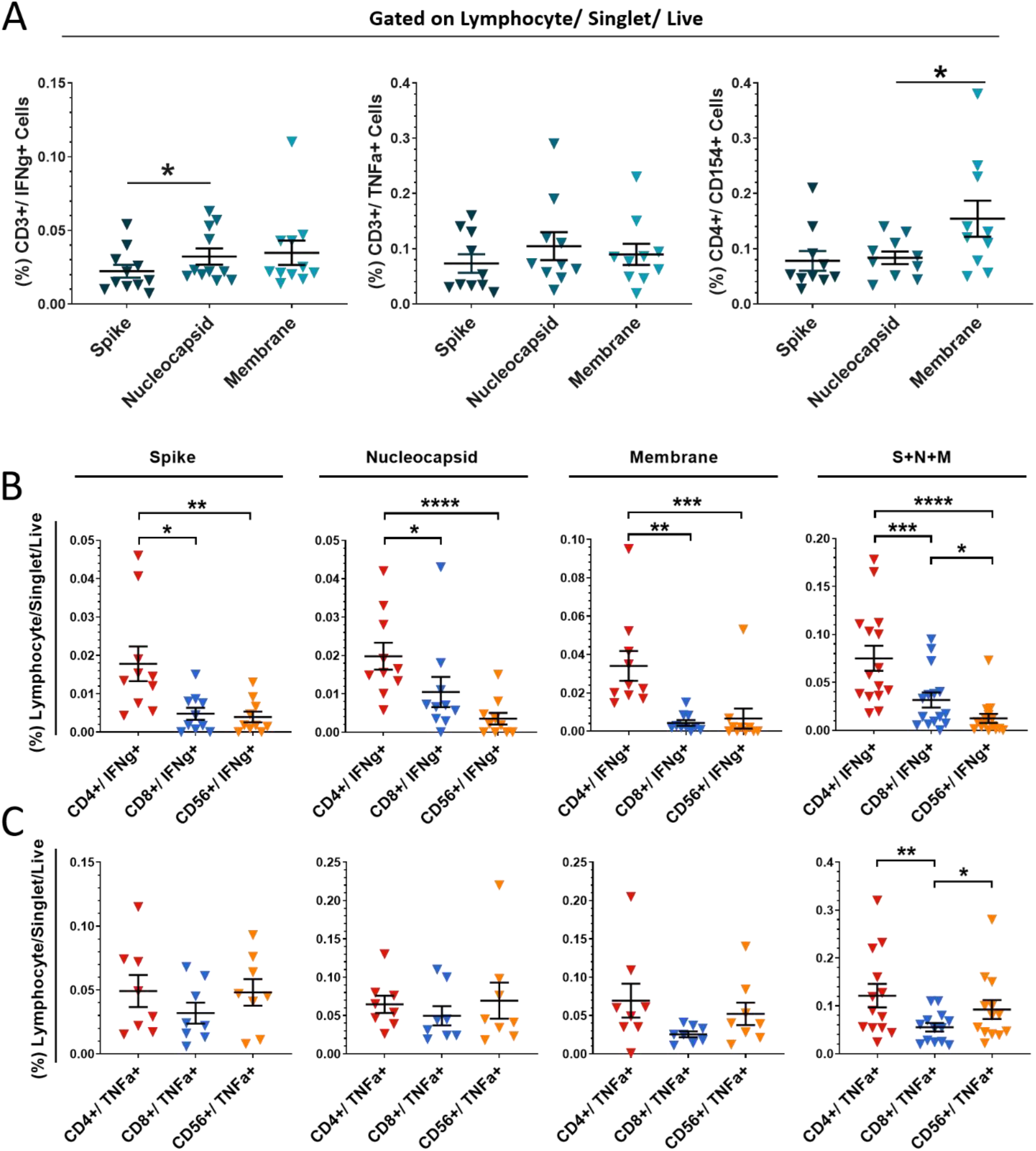
Covid-19 convalescent donor PBMC responses to SARS-CoV-2 peptides. **(A)** Mean percentages of CD3+/IFN-γ+ cells, CD3+/TNF-α+ cells and CD4+/CD154+ cells were compared for response to individual peptide pools: Spike, Nucleocapsid and Membrane in CCD (n=10). Data represented as mean ± SEM. Lymphocyte subsets (CD4+ T cell, CD8+ T cell and CD56+ NK cells) were compared for **(B)** IFN-γ response and **(C)** TNF-α response stimulation with individual peptide pools and combined SARS-CoV-2 peptide pools. Data represented as mean ± SEM. All significance was determined using RM one-way ANOVA with Geisser-Greenhouse correction where *p≤0.05, **p≤0.01, p≤0.001 and **** p≤ 0.0001.

**Figure S4.**
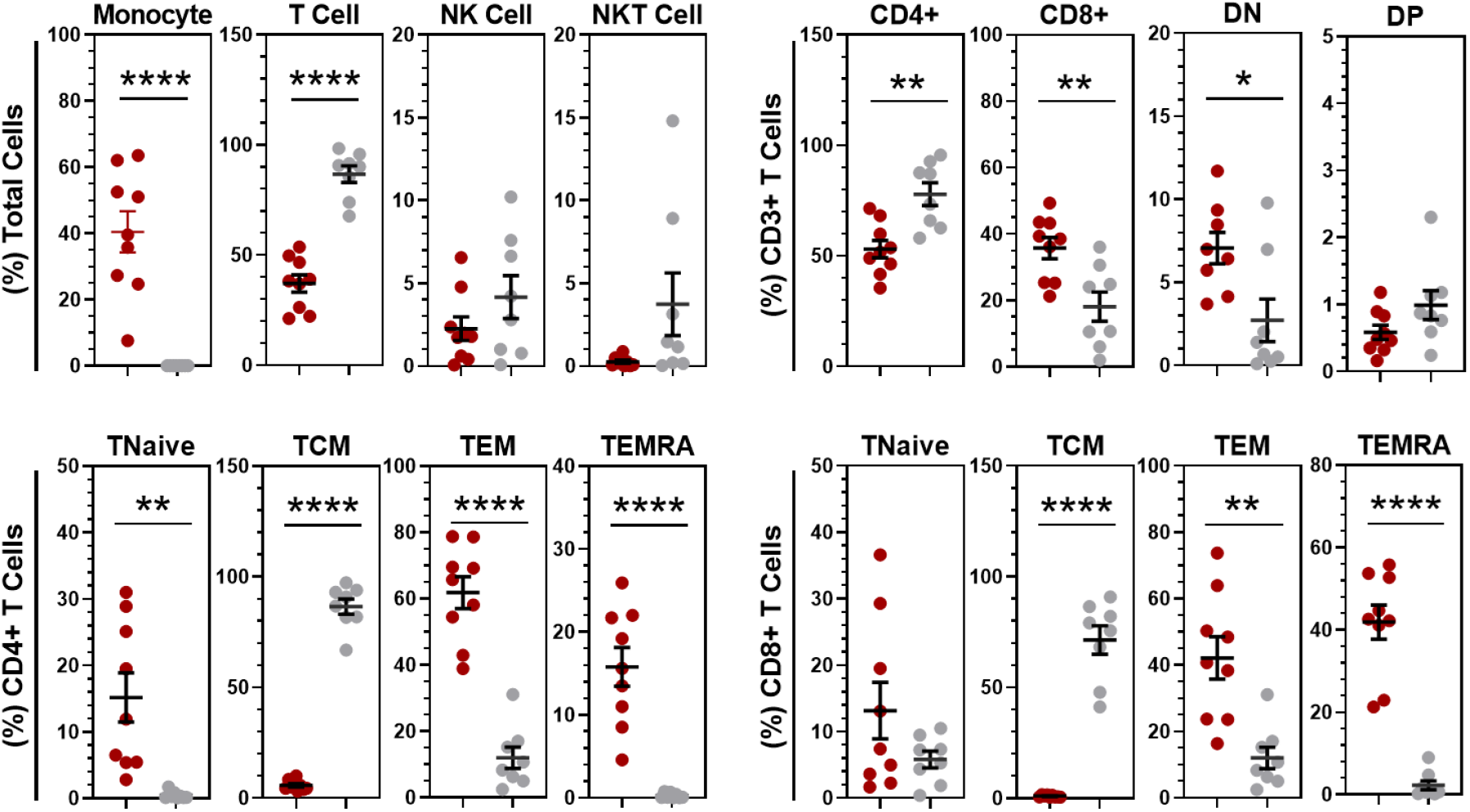
Direct comparison of T cell subpopulations between day 0 isolated VST and day 14 expanded VST. Significance was calculated using paired t-tests with Holm-Sidak correction for multiple comparisons *p≤0.05, **p≤0.01, p≤0.001 and **** p≤ 0.0001. All data is represented as mean ± SEM.

**Figure S5.**
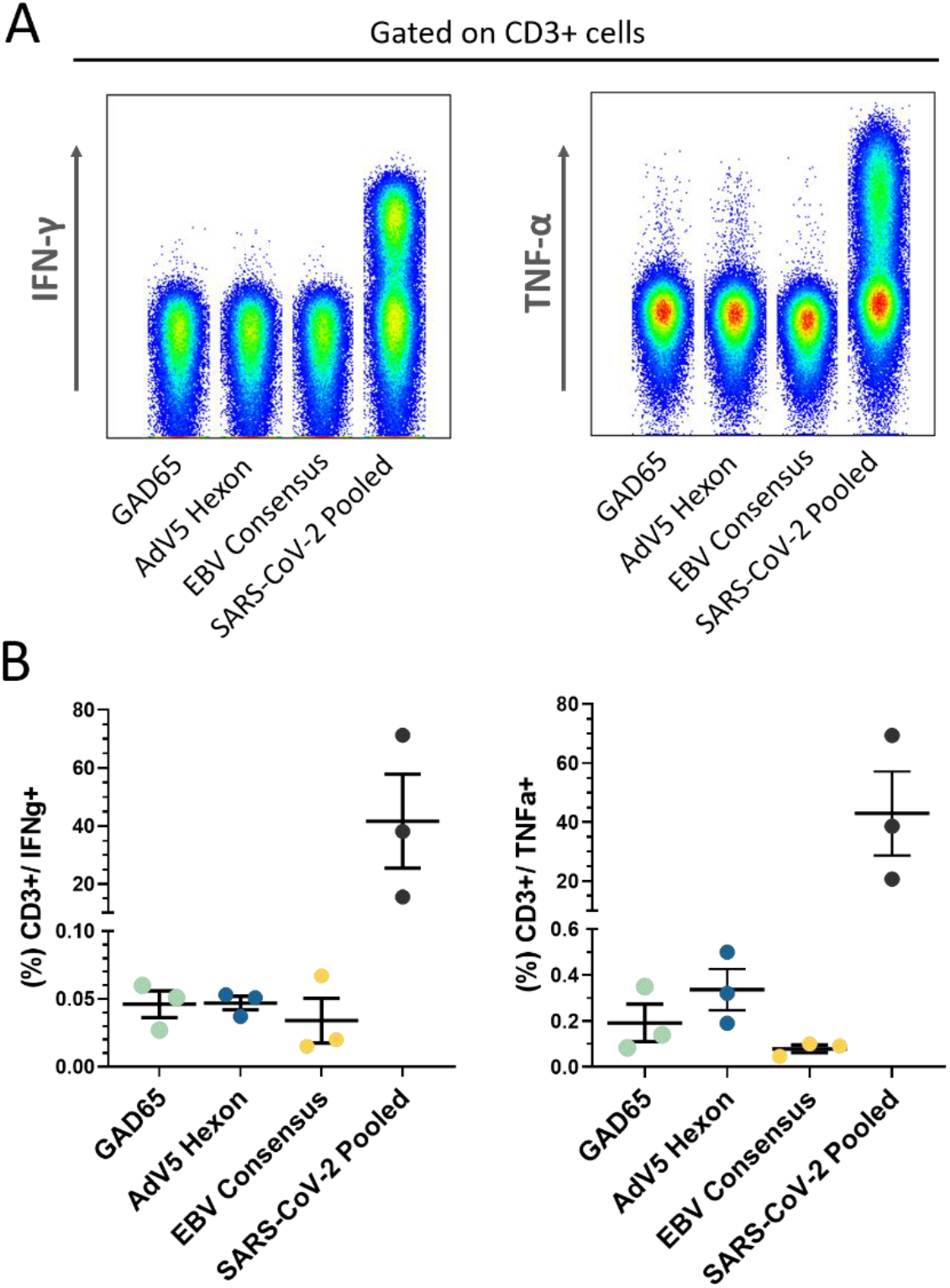
Specificity of cultured SARS-CoV-2 VSTs. SARS-CoV-2. VST cultures at day 14 were co-cultured with antigen-loaded mature autologous DCs and assessed for response. **(A)** Peptide specificity in a representative individual VST culture was assessed using DC loaded with GAD65 peptide, Adenovirus5 (AdV5) Hexon peptide, Epstein-Barr Virus (EBV) consensus peptide and combined pools of SARS-CoV-2 peptides (Spike + Nucleocapsid + Membrane) as positive control. **(B)** The mean percentage ± SEM of CD3+/IFN-γ+ cells, CD3+/TNF-α+ cells in day 14 SARS-CoV-2 VSTs (n=3) for each peptide. No statistical significance was determined.

## References

1 P. Zhou, X. L. Yang, X. G. Wang, B. Hu, L. Zhang, H. R. Si, B. Li, C. L. Huang, H. D. Chen, J. Chen, Y. Luo, H. Guo, R. D. Jiang, M. Q. Liu, Y. Chen, X. R. Shen, X. Wang, X. S. Zheng, K. Zhao, Q. J. Chen, F. Deng, L. L. Liu, B. Yan, F. X. Zhan, Y. Y. Wang, G. F. Xiao, Z. L. Shi, A pneumonia outbreak associated with a new coronavirus of probable bat origin. Nature. 579, 270–273 (2020).

2 F. Wu, S. Zhao, B. Yu, Y. M. Chen, W. Wang, Z. G. Song, Y. Hu, Z. W. Tao, J. H. Tian, Y. Y. Pei, M. L. Yuan, Y. L. Zhang, F. H. Dai, Y. Liu, Q. M. Wang, J. J. Zheng, L. Xu, E. C. Holmes, Y. Z. Zhang, A new coronavirus associated with human respiratory disease in China. Nature. 579, 265–269 (2020).

3 COVID-19 situation update worldwide, as of 30^th^ July 2020 (European Centre for Disease Prevention and Control, Situation updates on COVID-19). https://www.ecdc.europa.eu/en/geographical-distribution-2019-ncov-cases.

4 C. Huang, Y. Wang, X. Li, L. Ren, J. Zhao, Y. Hu, L. Zhang, G. Fan, J. Xu, X. Gu, Z. Cheng, T. Yu, J. Xia, Y. Wei, W. Wu, X. Xie, W. Yin, H. Li, M. Liu, Y. Xiao, H. Gao, L. Guo, J. Xie, G. Wang, R. Jiang, Z. Gao, Q. Jin, J. Wang, B. Cao, Clinical features of patients infected with 2019 novel coronavirus in Wuhan, China. Lancet. 395, 497–506 (2020).

5 D. Wang, B. Hu, C. Hu, F. Zhu, X. Liu, J. Zhang, B. Wang, H. Xiang, Z. Cheng, Y. Xiong, Y. Zhao, Y. Li, X. Wang, Z. Peng, Clinical Characteristics of 138 Hospitalized Patients with 2019 Novel Coronavirus-Infected Pneumonia in Wuhan, JAMA. 323, 1061–1069 (2020).

6 G. Grasselli, A. Zangrillo, A. Zanella, M. Antonelli, L. Cabrini, A. Castelli, D. Cereda, A. Coluccello, G. Foti, R. Fumagalli, G. lotti, N. Latronico, L. Lorini, S. Merler, G. Natalini, A. Piatti, M. V. Ranieri, A. M. Scandroglio, E. Storti, M. Cecconi, A. Pesenti, for the COVID-19 Lombardy ICU Network, Baseline Characteristics and Outcomes of 1591 Patients Infected with SARS-CoV-2 Admitted to ICUs of the Lombardy Region, Italy. JAMA. 323, 1574–1581 (2020).

7 J. Grein, N. Ohmagari, D. Shin, G. Diaz, E. Asperges, A. Castagna, T. Feldt, G. Green, M. L. Green, F. X. Lescure, E. Nicastri, R. Oda, K. Yo, E. Quiros-Roldan, A. Studemeister, J. Redinski, S. Ahmed, J. Bernett, D. Chelliah, D. Chen, S. Chihara, S. H. Cohen, J. Cunningham, A. D’Arminio Monforte, S. Ismail, H. Kato, G. Lapadula, E. L’Her, T. Maeno, S. Majumder, M. Massari, M. Mora-Rillo, Y. Mutoh, D. Nguyen, E. Verweij, A. Zoufaly, A. O. Osinusi, A. DeZure, Y. Zhao, L. Zhong, A. Chokkalingam, E. Elboudwarej, L. Telep, L. Timbs, I. Henne, S. Sellers, H. Cao, S. K. Tan, L. Winterbourne, P. Desai, R. Mera, A. Gaggar, R. P. Myers, D. M. Brainard, R. Childs, T. Flanigan, Compassionate Use of Remdesivir for Patients with Severe COVID-19. N Engl J Med. 382, 2327–2336 (2020).

8 P. Horby, W. S. Lim, J. R. Emberson, M. Mafham, J. L. Bell, L. Linsell, N. Staplin, C. Brightling, A. Ustianowski, E. Elmahi, B. Prudon, C. Green, T. Felton, D. Chadwick, K. Rege, C. Fegan, L. C. Chappell, S. N. Faust, T. Jaki, K. Jeffery, A. Montgomery, K. Rowan, E. Juszczak, J. K. Baillie, R. Haynes, M. J. Landray, Dexamethasone in Hospitalized Patients with COVID-19: Preliminary Report. N Engl J Med. Online ahead of print. https://www.nejm.org/doi/10.1056/NEJMoa2021436

9 https://www.synairgen.com/. Accessed 23rd July 2020.

10 M. Zheng, Y. Gao, G. Wang, G. Song, S. Liu, D. Sun, Y. Xu, Z. Tian, Functional exhaustion of antiviral lymphocytes in COVID-19 patients. Cell Mol Immunol. 17, 533–535 (2020).

11 C. Qin, L. Zhou, Z. Hu, S. Zhang, S. Yang, Y. Tao, C. Xie, K. Ma, K. Shang, W. Wang, D. S. Tian, Dysregulation of immune response in patients with COVID-19 in Wuhan, China. Clin Infect Dis. 71, 762–768 (2020).

12 L. F. Garcia; Immune Response, Inflammation, and the Clinical Spectrum of COVID-19. Front Immunol, 16 June 2020. https://doi.org/10.3389/fimmu.2020.01441

13 M. Z. Tay, C. M. Poh, L. Rénia, P. A. MacAry, L. F. P. Ng, The trinity of COVID-19: immunity, inflammation and intervention. Nat Rev Immunol. 20, 363–74 (2020). doi: 10.1038/s41577-020-0311-8

14 X. Cao, COVID-19: immunopathology and its implications for therapy. Nat Rev Immunol. 20, 269–70 (2020). doi: 10.1038/s41577-020-0308-3

15 R. H. Manjili, M. Zarei, M. Habibi, M. H. Manjili, COVID-19 as an acute inflammatory disease. J Immunol. 205, 12–19 (2020). doi: 10.4049/jimmunol.2000413

16 S. Felsenstein, J. A. Herbert, P. S. McNamara, C. M. Hedrich, COVID-19: Immunology and treatment options. Clin Immunol. 215, 108448 (2020). doi: 10.1016/j.clim.2020.108448

17 J. D. M. Campbell, A. Foerster, V. Lasmanowicz, M. Niemöller, A. Scheffold, M. Fahrendorff, G. Rauser, M. Assenmacher, A. Richter; Rapid detection, enrichment and propagation of specific T cell subsets based on cytokine secretion. Clin Exp Immunol. 163, 1–10 (2011).

18 G. Rauser, H. Einsele, C. Sinzger, D. Wernet, G. Kuntz, M. Assenmacher, J. D. M. Campbell, M. S. Topp, Rapid generation of combined CMV-specific CD4+ and CD8+ T-cell lines for adoptive transfer into recipients of allogeneic stem cell transplants. Blood. 103, 3565–3572 (2004).

19 T. Feuchtinger, S. Matthes-Martin, C. Richard, T. Lion, M. Fuhrer, K. Hamprecht, R. Handgretinger, C. Peters, F. R. Schuster, R. Beck, M. Schumm, R. Lotfi, G. Jahn, P. Lang, Safe adoptive transfer of virus-specific T-cell immunity for the treatment of systemic adenovirus infection after allogeneic stem cell transplantation. Br J Haematol. 134, 64–76 (2006).

20 S. Kazi, A. Mathur, G. Wilkie, K. Cheal, R. Battle, N. McGowan, N. Fraser, E. Hargreaves, D. Turner, J. D. M. Campbell, M. Turner, M. A. Vickers, Long-term follow up after third-party viral-specific cytotoxic lymphocytes for immunosuppression- and Epstein-Barr virus-associated lymphoproliferative disease. Haematologica. 104, e356–e359 (2019).

21 I. Tzannou, A. Watanabe, S. Naik, R. Daum, M. Kuvalekar, K. S. Leung, C. Martinez, G. Sasa, M. Wu, A. P. Gee, R. A. Krance, S. Gottschalk, H. E. Heslop, B. Omer, “Mini” bank of only 8 donors supplies CMV-directed T cells to diverse recipients. Blood Adv. 3, 2571–2580 (2019).

22 T. Haque, G. M. Wilkie, M. M. Jones, C. D. Higgins, G. Urquhart, P. Wingate, D. Burns, K. McAulay, M. Turner, C. Bellamy, P. L. Amlot, D. Kelly, A. MacGilchrist, M. K. Gandhi, A. J. Swerdlow, D. H. Crawford, Allogeneic cytotoxic T-cell therapy for EBV positive posttransplantation lymphoproliferative disease: results of a phase 2 multicenter clinical trial. Blood. 110, 1123–1131 (2007).

23 J. Liu, S. Li, J. Liu, B. Liang, X. Wang, H. Wang, W. Li, Q. Tong, J. Yi, L. Zhao, L. Xiong, C. Guo, J. Tian, J. Luo, J. Yao, R. Pang, H. Shen, C. Peng, T. Liu, Q. Zhang, J. Wu, L. Xu, S. Lu, B. Wang, Z. Weng, C. Han, H. Zhu, R. Zhou, H. Zhou, X. Chen, P. Ye, B. Zhu, L. Wang, W. Zhou, S. He, Y. He, S. Jie, P. Wei, J. Zhang, Y. Lu, W. Wang, L. Zhang, L. Li, F. Zhou, J. Wang, U. Dittmer, M. Lu, Y. Hu, D. Yang, X. Zheng, Longitudinal characteristics of lymphocyte responses and cytokine profiles in the peripheral blood of SARS-CoV-2 infected patients. EBioMedicine. May 2020 102763. doi: 10.1016/j.ebiom.2020.102763.

24 J. W. Song, C. Zhang, X. Fan, F. P. Meng, Z. Xu, P. Xia, W. J. Cao, T. Yang, X. P. Dai, S. Y. Wang, R. N. Xu, T. J. Jiang, W. G. Li, D. W. Zhang, P. Zhao, M. Shi, C. Agrati, G. Ippolito, M. Maeurer, A. Zumla, F. S. Wang, J. Y. Zhang, Immunological and inflammatory profiles in mild and severe cases of COVID-19. Nat Commun. 11, 3410 (2020). doi: 10.1038/s41467-020-17240-2

25 G. Chen, D. Wu, W. Guo, Y. Cao, D. Huang, H. Wang, T. Wang, X. Zhang, H. Chen, H. Yu, X. Zhang, M. Zhang, S. Wu, J. Song, T. Chen, M. Han, S. Li, X. Luo, J. Zhao, Q. Ning, Clinical and immunological features of severe and moderate coronavirus disease 2019. J Clin Invest. 130, 2620–2629 (2020). doi: 10.1172/JCI137244.

26 S. De Biasi, M. Meschiari, L. Gibellini, C. Bellinazzi, R. Borella, L. Fidanza, L. Gozzi, A. Iannone, D. Lo Tartaro, M. Mattioli, A. Paolini, M. Menozzi, J. Milić, G. Franceschi, R. Fantini, R. Tonelli, M. Sita, M. Sarti, T. Trenti, L. Brugioni, L. Cicchetti, F. Facchinetti, A. Pietrangelo, E. Clini, M. Girardis, G. Guaraldi, C. Mussini, A. Cossarizza, Marked T cell activation, senescence, exhaustion and skewing towards TH17 in patients with COVID-19 pneumonia. Nat Commun. 11, 3434 (2020). doi: 10.1038/s41467-020-17292-4.

27 T. Kaeuferle, R. Krauss, F. Blaeschke, S. Willier, T. Feuchtinger, Strategies of adoptive T -cell transfer to treat refractory viral infections post allogeneic stem cell transplantation. J Hematol Oncol. 12, 13 (2019).

28 T. Sekine, A. Perez-Potti, O. Rivera-Ballesteros, K. Strålin, J. B. Gorin, A. Olsson, S. Llewellyn-Lacey, H. Kamal, G. Bogdanovic, S. Muschiol, D. J. Wullimann, T. Kammann, Emgård, T. Parrot, E. Folkesson, O. Rooyackers, L. I. Eriksson, A. Sönnerborg, T. Allander, J. Albert, M. Nielsen, J. Klingström, S. Gredmark-Russ, N. K. Björkström, J. Sandberg, D. A. Price, H. G. Ljunggren, S. Aleman, M. Buggert, Robust T cell immunity in convalescent individuals with asymptomatic or mild COVID-19. BioRxiv preprint doi: https://doi.org/10.1101/2020.06.29.174888

29 T. P. Plackett, E. D. Boehmer, D. E. Faunce, E. J. Kovacs, Aging and innate immune cells. J Leukoc Biol. 76, 291–299 (2004).

30 N. H. Overgaard, J. W. Jung, R. J. Steptoe, J. W. Wells, CD4+/CD8+ double-positive T cells: more than just a developmental stage? J Leukoc Biol. 97, 31–38 (2015).

31 J. Braun, L. Loyal, M. Frentsch, D. Wendisch, P. Georg, F. Kurth, S. Hippenstiel, M. Dingeldey, B. Kruse, F. Fauchere, E. Baysal, M. Mangold, L. Henze, R. Lauster, M. A. Mall, K. Beyer, J. Röhmel, S. Voigt, J. Schmitz, S. Miltenyi, I. Demuth, M. A Müller, A. Hocke, M. Witzenrath, N. Suttorp, F. Kern, U. Reimer, H. Wenschuh, C. Drosten, V. M. Corman, C. Giesecke-Thiel, L. E. Sander, A. Thiel, SARS-CoV-2-reactive T cells in healthy donors and patients with COVID-19. Nature. July 2020 doi: 10.1038/s41586-020-2598-9.

32 A. Grifoni, D. Weiskopf, S. I. Ramirez, J. Mateus, J. M. Dan, C. Rydyznski Moderbacher, S. A. Rawlings, A. Sutherland, L. Premkumar, R. S. Jadi, D. Marrama, A. M. de Silva, A. Frazier, A. F. Carlin, J. A. Greenbaum, B. Peters, F. Krammer, D. M. Smith, S. Crotty, A. Sette, Targets of T Cell Responses to SARS-CoV-2 Coronavirus in Humans with COVID-19 Disease and Unexposed Individuals. Cell. 181, 1489–1501 (2020).

33 Y. Peng, A. J. Mentzer, G. Liu, X. Yao, Z. Yin, D. Dong, W. Dejnirattisai, T. Rostron, P. Supasa, C. Liu, C. Lopez-Camacho, J. Slon-Campos, Y. Zhao, D. Stuart, G. Paeson, J. Grimes, F. Antson, O. W. Bayfield, D. E. Hawkins, D. E. Ker, L. Turtle, K. Subramaniam, P. Thomson, P. Zhang, C. Dold, J. Ratcliff, P. Simmonds, T. de Silva, P. Sopp, D. Wellington, U. Rajapaksa, Y. L. Chen, M. Salio, G. Napolitani, W. Paes, P. Borrow, B. Kessler, J. W. Fry, N. F. Schwabe, M. G. Semple, K. J. Baillie, S. Moore, P. J. Openshaw, A. Ansari, S. Dunachie, E. Barnes, J. Frater, G. Kerr, P. Goulder, T. Lockett, R. Levin, R. J. Cornall, C. Conlon, P. Klenerman, A. McMichael, G. Screaton, J. Mongkolsapaya, J. C. Knight, G. Ogg, T. Dong, Broad and strong memory CD4+ and CD8+ T cells induced by SARS-CoV-2 in UK convalescent COVID-19 patients. BioRxiv. June 2020 https://doi.org/10.1101/2020.06.05.134551

34 A.J. Wilk, A. Rustagi, N. Q. Zhao, J. Roque, G. J. Martínez-Colón, J. L. McKechnie, G. T. Ivison, T. Ranganath, R. Vergara, T. Hollis, L. J. Simpson, P. Grant, A. Subramanian, A. J. Rogers, C. A. Blish, A single-cell atlas of the peripheral immune response in patients with severe COVID-19. Nat Med. 26, 1070–1076 (2020).

35 R. Channappanavar, C. Fett, J. Zhao, D. K. Meyerholx, S. Perlman, Virus-specific memory CD8 T cells provide substantial protection from lethal severe acute respiratory syndrome coronavirus infection. J Virol. 88, 11034–11044 (2014).

36 C. K. Kang, G. C. Han, M. Kim, G. Kim, H. M. Shin, K. H. Song, P. G. Choe, W. B. Park, E. S. Kim, H. B. Kim, N. J. Kim, H. R. Kim, M. D. Oh, Aberrant hyperactivation of cytotoxic T-cell as a potential determinant of COVID-19 severity. Int J Infect Dis. 97, 313–321 (2020).

37 A. Nguyen, J. K. David, S. K. Maden, M. A. Wood, B. R. Weeder, A. Nellore, R. F. Thompson. J Virol. 94, e00510–20 (2020).

38 D. Mathew, J. R. Giles, A. E. Baxter, A. R. Greenplate, J. E. Wu, C. Alanio, D. A. Oldridge, L. Kuri-Cervantes, M. Betina Pampena, K. D’Andrea, S. Manne, Z. Chen, Y. J. Huang, J. P. Reilly, A. R. Weisman, C. A. G. Ittner, O. Kuthuru, J. Dougherty, K. Nzingha, N. Han, J. Kim, A. Pattekar, E. C. Goodwin, E. M. Anderson, M. E. Weirick, S. Gouma, C. P. Arevalo, M. J. Bolton, F. Chen, S. F. Lacey, S. E. Hensley, S. Apostolidis, A. C. Huang, L. A. Vella, UPenn COVID Processing Unit; M. R. Betts, N. J. Meyer, E. J. Wherry, Deep immune profiling of COVID-19 patients reveals distinct immunotypes with therapeutic implications. Science. July 2020 doi: 10.1126/science.abc8511

